# *De Novo* Prediction of Human Chromosome Structures: Epigenetic Marking Patterns Encode Genome Architecture

**DOI:** 10.1101/173088

**Authors:** Michele Di Pierro, Ryan R. Cheng, Erez Lieberman Aiden, Peter G. Wolynes, José N. Onuchic

## Abstract

Inside the cell nucleus, genomes fold into organized structures that are characteristic of cell type. Here, we show that this chromatin architecture can be predicted *de novo* using epigenetic data derived from ChIP-Seq. We exploit the idea that chromosomes encode a one-dimensional sequence of chromatin structural types. Interactions between these chromatin types determine the three-dimensional (3D) structural ensemble of chromosomes through a process similar to phase separation. First, a recurrent neural network is used to infer the relation between the epigenetic marks present at a locus, as assayed by ChIP-Seq, and the genomic compartment in which those loci reside, as measured by DNA-DNA proximity ligation (Hi-C). Next, types inferred from this neural network are used as an input to an energy landscape model for chromatin organization (MiChroM) in order to generate an ensemble of 3D chromosome conformations. After training the model, dubbed MEGABASE (Maximum Entropy Genomic Annotation from Biomarkers Associated to Structural Ensembles), on odd numbered chromosomes, we predict the chromatin type sequences and the subsequent 3D conformational ensembles for the even chromosomes. We validate these structural ensembles by using ChIP-Seq tracks alone to predict Hi-C maps as well as distances measured using 3D FISH experiments. Both sets of experiments support the hypothesis of phase separation being the driving process behind compartmentalization. These findings strongly suggest that epigenetic marking patterns encode sufficient information to determine the global architecture of chromosomes and that *de novo* structure prediction for whole genomes may be increasingly possible.

## Main Text

In the nucleus of eukaryotic cells, the one-dimensional information of the genome is organized in three dimensions (1, 2). It is increasingly evident that genomic spatial organization is a key element of transcriptional regulation (1, 3). During interphase, the three-dimensional (3D) arrangement of chromatin brings into close spatial proximity sections of DNA separated by great genomic distance, introducing interactions between genes and regulatory elements. These folding patterns are cell-type specific (4, 5), and their disruption can lead to disease (6-9).

The use of high-resolution contact mapping experiments (Hi-C) has revealed that, at the large scale, genome structure is dominated by the segregation of human chromatin into compartments. Initial analysis of Hi-C experiments revealed that loci typically exhibited one of two long-range contact patterns, suggesting the presence of two spatial neighborhoods, dubbed the A and B compartments. (10). Subsequently, higher resolution experiments have shown the presence of six distinct long-range patterns, indicating the presence of six sub-compartments (A1, A2, B1, B2, B3, and B4) in human lynphoblastoid cells (GM12878)(5). The compartmentalization of the genome has been observed in many organisms (including mouse (5, 11) and *Drososphila* (12, 13)), and has been confirmed by numerous microscopy experiments (14). Crucially, the long-range contact pattern seen at a locus is cell-type specific, and is strongly associated with particular chromatin marks.

To model this structure, we recently introduced an effective energy landscape model for chromatin structure called the Minimal Chromatin Model (MiChroM) (15). This model combines a generic polymer potential with additional interaction terms governing compartment formation as well as other processes involved in chromatin organization— i.e., the local helical structural tendency of the chromatin filament (15-18) and the chromatin loops associated with the presence of CCCTC-binding factor (CTCF) (5, 19-21). The formation of compartments (as well as any other interaction in MiChroM) is assumed to operate only through direct protein-mediated contacts, bringing about segregation of chromatin types through a process of phase separation (15, 22). MiChroM shows that the compartmentalization patterns that Hi-C maps reveal can be transformed into 3D models of genome structure at 50 kb resolution.

Here, we extend the earlier work by demonstrating that the structure of whole genomes can be predicted, *de novo*, by inferring chromatin types from ChIP-Seq data and then using these inferences as an input into an effective energy landscape model. The workflow behind this approach is broadly described in Figure 1.

**Figure 1.**
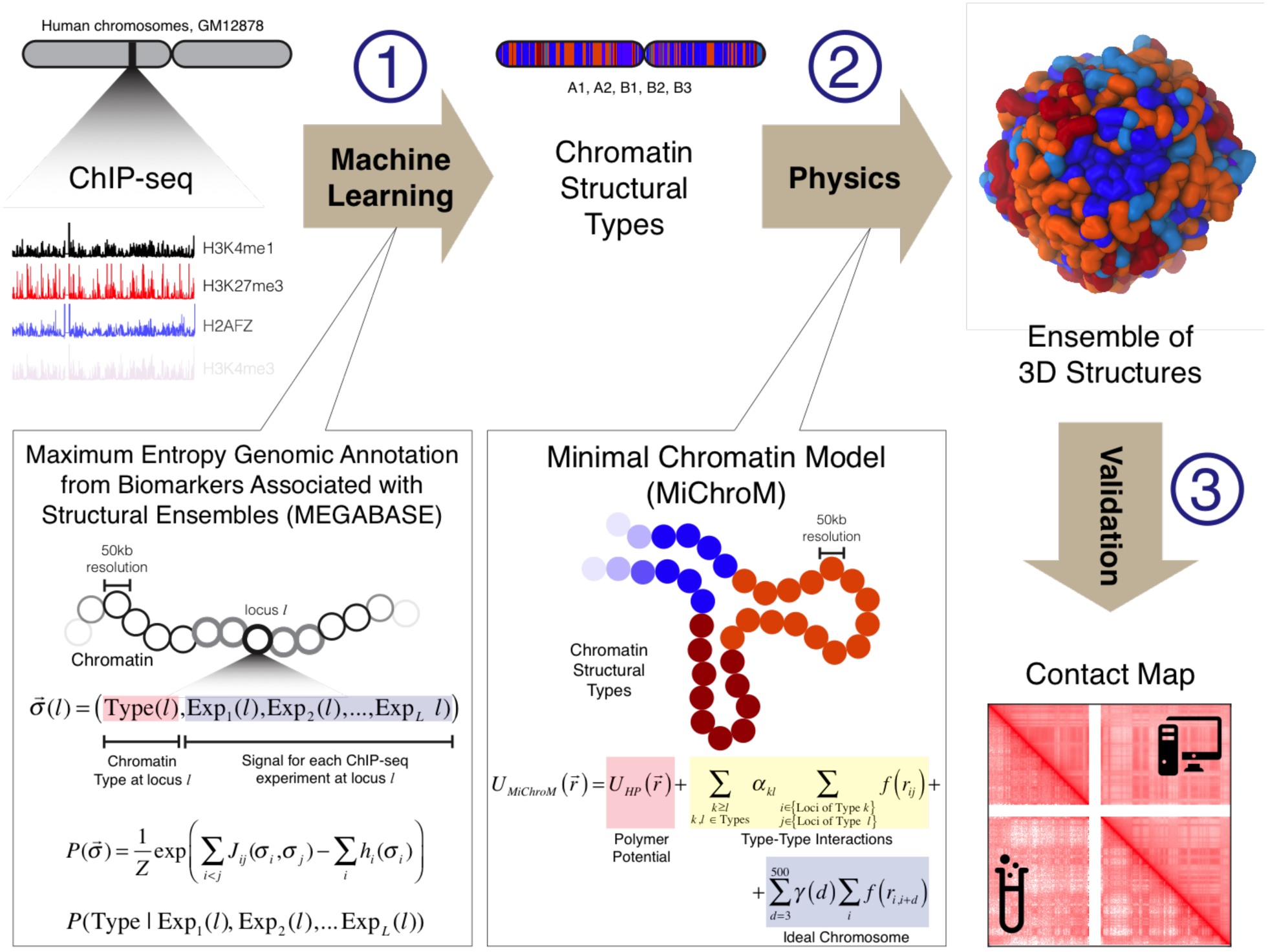
Schematic illustration of the MEGABASE+MiChroM computational pipeline. **(1) ChIP-seq data constitutes the only input to our pipeline**. ChIP-Seq tracks obtained from a publicly available resource (ENCODE) are converted into a sequence of chromatin structural types using a recurrent neural network dubbed MEGABASE. The neural network encodes the relationship between compartmentalization and the biochemical state of each locus along the genome. **(2) Sequences of chromatin structural types are used as input to a physical model for chromatin folding (MiChroM) to obtain the ensembles of 3D structures of specific chromosomes** (15). MiChroM is an effective energy landscape model consisting of a generic polymer with chromatin type interactions and a translational invariant local ordering term (Ideal Chromosome). **(3) Ensembles of 3D structures are validated by comparing the predicted contact maps with those experimentally determined by using Hi-C.**

Although the compartments and sub-compartments visible in Hi-C maps correlate with a handful of specific epigenetic modifications present at those loci (see also (5)), the distributions of epigenetic markers found in each compartment are broad and largely overlap. It is therefore impossible to assign correctly any given locus to a specific compartment using the frequency of any single epigenetic modification. To overcome this difficulty, we use a machine learning approach to extract information from the raw chromatin immunoprecipitation (ChIP-Seq) data. We first obtained ChIP-Seq profiles available from the ENCODE project for the GM12878 lymphoblastoid cell line, encompassing 84 protein-binding experiments and 11 histone marks. Next, we discretized each of these profiles, partitioning them into 50 kb loci, each of which is assigned a value from 1 (weakest signal) to 20 (strongest signal). We then constructed a recurrent neural network to uncover the relationship between compartment annotations and epigenetic markings. We use a neural network in which each data type available at a given locus corresponds to a single neuron (23). The state of the network is represented by the state vector *σ⃗*(*l*) = (*C*(*l*), Exp_1_(*l*),Exp_2_(*l*), …, Exp_*L*_(*l*)), which represents all the data available at locus *l*—with *C* being the sub-compartment annotation and Exp_*i*_. being the result of the *i*-th ChIP-Seq experiment. The data at each locus are further assumed to be distributed according to a Boltzmann Distribution for a Potts Model:

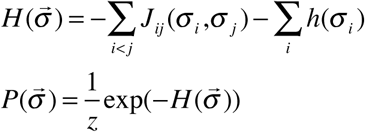

where the *P*(*σ⃗*) indicates the probability of observing the state vector *σ⃗* at any given locus *l*; the *J_ij_* interactions capture local pairwise correlations between epigenetic marks or between marks and chromatin types; and the *h*’s determine the individual frequencies of chromatin types and markers. This procedure is equivalent to training a Boltzmann Machine to encode the information contained in the data set. The learning strategy is based on the idea that the parameters of the neural network should maximize the likelihood of observing the set of state vectors representing a particular training set. A similar strategy has been previously introduced to quantify the correlated mutational patterns observed in amino acid sequence data of protein families occurring under natural selection to aid protein structure prediction (24, 25).

The quality of compartment prediction is improved when we include in the Potts Model interactions that do not just refer to a single 50 kb locus but also interactions encoding correlations between markings and annotations of nearest neighbors and next nearest neighbors (i.e. the neural network correlates information from loci *l*-2, *l*-1, *l*, *l*+*1*, *l*+2). Through these couplings, the probability of observing a specific state vector at a given locus is correlated with the states of the adjacent segments thus minimizing the effect of uncorrelated noise. This strategy is analogous to the construction of secondary structure predictors in protein folding using helix-coil models (26).

The inferred probabilistic model is then marginalized to predict the most probable chromatin type for a given locus *l* when given the experimental ChIP-Seq measurements of loci (*l* – 2, *l* – 1, *l*, *l* +1, *l* + 2):

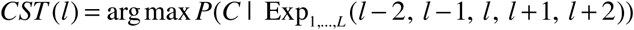

We refer to the resulting probabilistic predictor of chromatin structural types as the Maximum Entropy Genomic Annotation from Biomarkers Associated to Structural Ensembles (MEGABASE). Once trained for a given new input sequence of epigenetic marks the model can then find the most probable sequence of corresponding compartment annotations.

The state vectors of every locus of the odd numbered chromosomes comprise the training set. The state vectors of the even numbered chromosomes then provide a test set to quantify the performance of the trained model.

After training on the odd numbered chromosomes, we used our statistical model to predict the chromatin types for the independent set of the even chromosomes of the cell line GM12878 from their epigenetic marking profiles. For the test set, the predicted type assignments are in broad agreement with the experimentally determined structural annotations in (5). Specifically, the model is very accurate in predicting the assignments to compartments (A vs. B) while producing a larger number of mismatches between the predicted chromatin types and the published subcompartments annotations, which are more fine-grained (A1 vs. A2, B1 vs. B2 vs. B3). (See Figure S1)

Once predicted sequences of type annotations are available, we use our earlier MiChroM model to sample the predicted conformational ensembles of three-dimensional structures. In order to highlight the relationship between chromatin types and compartmentalization, we use the MiChroM Hamiltonian with the same parameters that had already been determined but omit the term in that energy function that models the CTCF-mediated looping interactions. These looping interactions seem to arise from a distinct process from compartmentalization and omitting such interactions does not disrupt the large-scale architecture of chromosomes (15) (See Figure S2 for the results of additional simulations including also the CTCF-mediated looping interactions).

The simulations all start from a random collapsed polymer having the proper length confined in a spherical region at correct density (See S.I.). After equilibration, we collect an ensemble of 3D structures representing the chromosome-specific energy landscape as shaped by the input inferred chromatin type sequence and by the MiChroM effective interactions.

From the ensemble of equilibrium conformations we calculate the contact probabilities between any pair of loci within each chromosome. We compare the resulting contact maps from the simulated ensemble of 3D structures with the experimental Hi-C maps from ref. (5). The overall agreement between the experimental and simulated contact probabilities is visually evident. The comparison between the simulated and experimental contact maps is shown in Figure 2 for representative chromosomes in the test set (i.e., the even autosomes). The Pearson’s coefficient is ~ 0.9 or higher for all of the chromosomes whether in the training set or test set and the analysis of the Pearson’s coefficient as a function of genomic distance (Figure S3-24) confirms that the two sets of maps are exceptionally well correlated. The power law scaling of the contact probability between two loci as a function of their genomic distance is well-reproduced at all genomic distances when comparing with Hi-C data (Figure S3-24).

**Figure 2.**
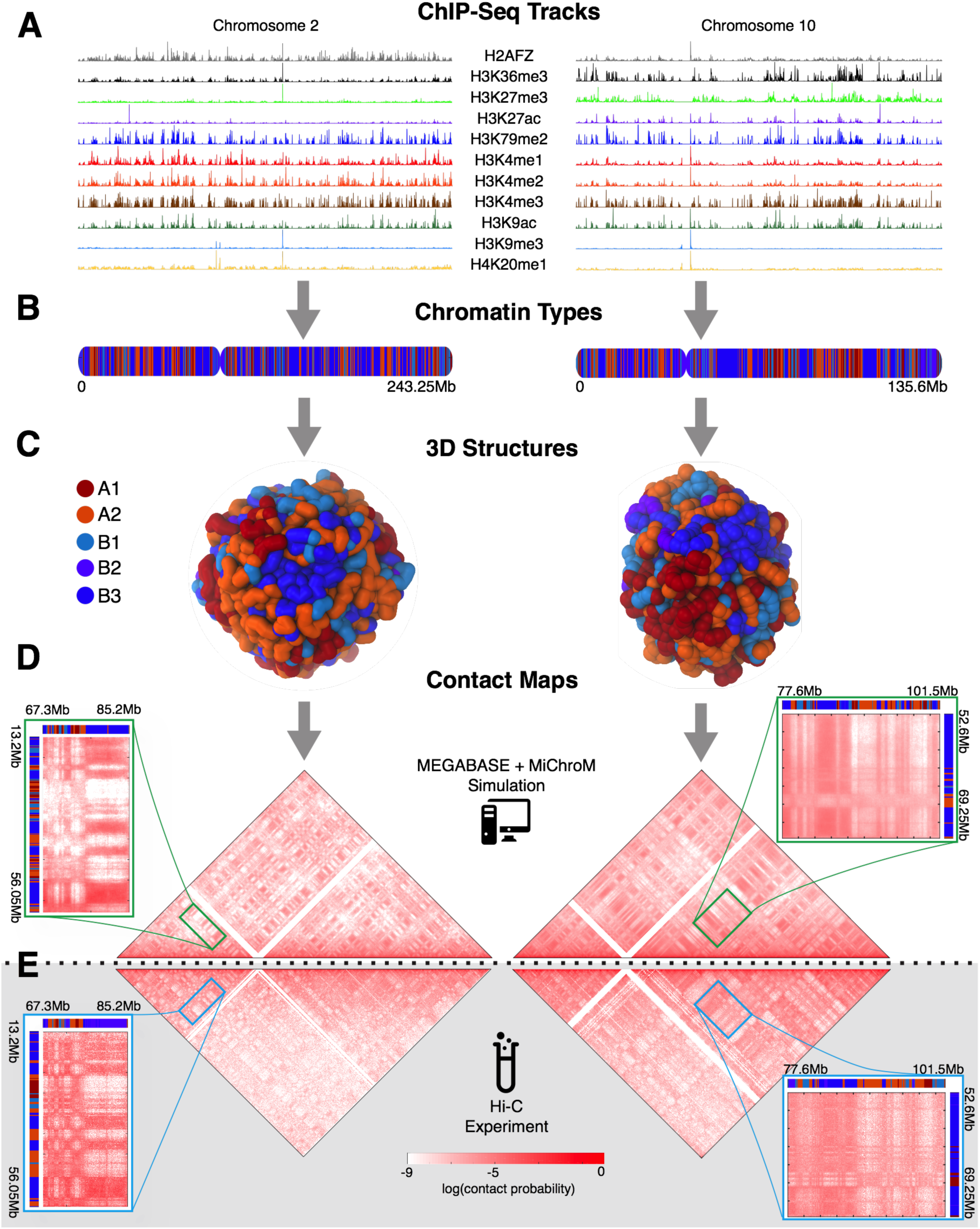
Predicting the 1D chromatin sequences, 3D conformations, and 2D contact probabilities of human chromosomes from epigenetic marking patterns. We apply MEGABASE + MiChroM to obtain an ensemble of 3D structures for all of the autosomes of cell line GM12878. For illustrative purposes, predictions for chromosomes 2 and 10 are shown on the left and right columns, respectively. (A) 95 ChIP-Seq tracks are downloaded from ENCODE and used as input for MEGABASE to predict 1D sequences of chromatin types shown in (B). The 3D structure of each chromosome is encoded in its specific 1D sequence of chromatin structural types. A typical 3D conformation obtained by MiChroM is shown in (C) for chromosome 2 and 10. Approximately 50,000 structures are collected from simulation to generate high-quality contact maps shown in (D). These contact maps are compared with the Hi-C maps shown in (E). The simulations correctly predict the long-range contact probability patterns that are observed in Hi-C maps, as seen in the magnified regions.

Finally, we compare the Cartesian distances between multiple pairs of loci as predicted through the use of our computational model with those measured by using 3D Fluorescence In Situ Hybridization (FISH), and reported in ref. (5) for the cell line GM12878 and in ref. (10) for the closely related cell line GM06990. FISH experiments in Figure 3 show that chromatin belonging to the same structural type tends to come into contact more frequently than otherwise, supporting the idea that compartmentalization is induced by a process of phase separation. This behavior is predicted with quantitative accuracy by our ChIP-Seq based simulation. Remarkably, simulations predict all the experimentally determined average distances, together with their variances (Figure 3, Figure S25, and Figure S26).

**Figure 3.**
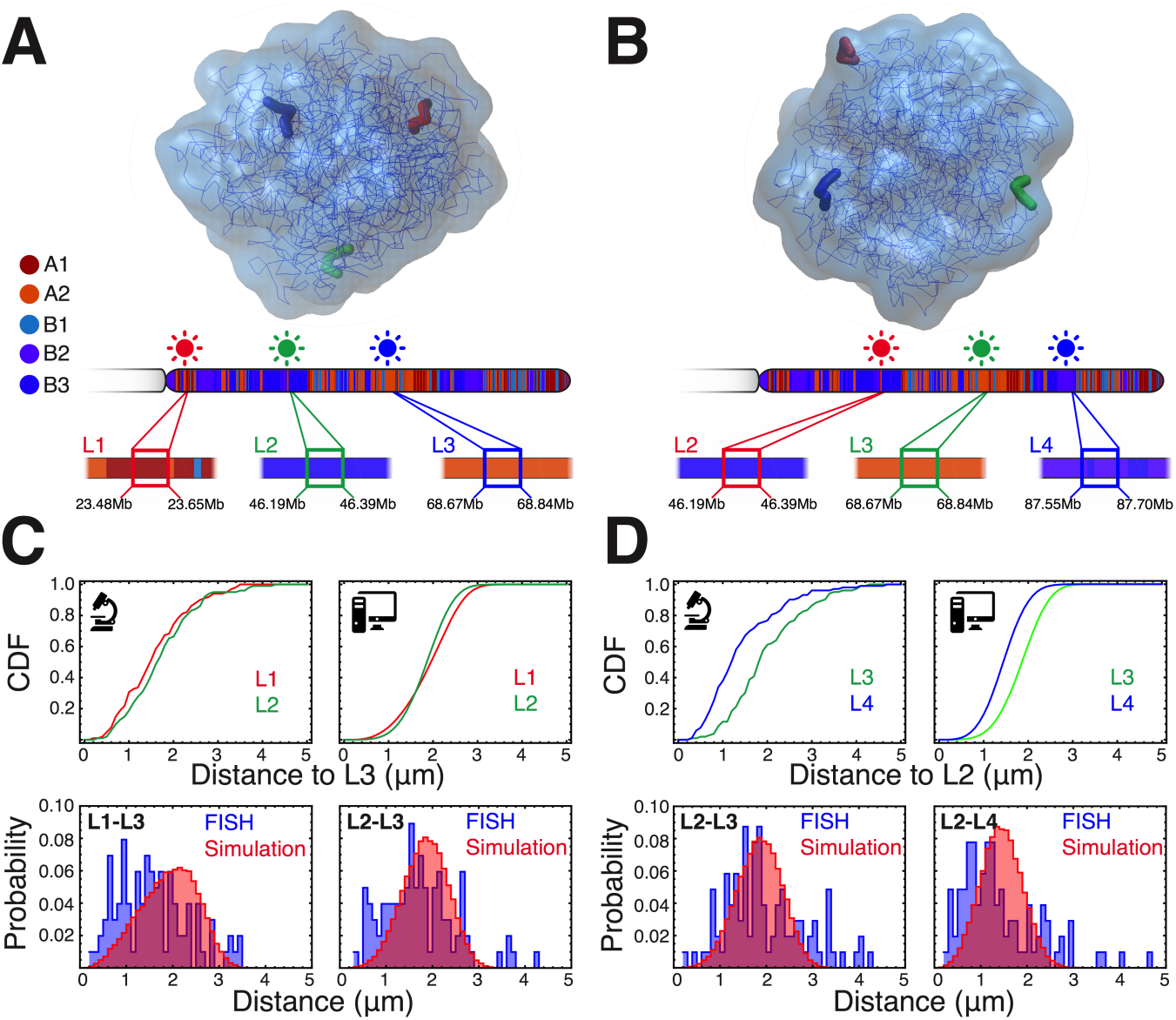
Simulated conformational ensembles predict the distances measured by 3D FISH experiments. Simulations and 3D Fluorescence In Situ Hybridization (FISH) experiments support the idea that compartmentalization observed in Hi-C maps emerges from the phase separation of chromatin structural types. (A and B) The Cartesian distances between four loci (L1, L2, L3, and L4) in chromosome 14 (cell line GM06990) were measured in two distinct 3D FISH experiments reported in ref. (10). The same distances were measured using the MEGABASE+MiChroM pipeline^1^. The positions of the fluorescent probes are illustrated in representative 3D configurations from simulations as well as along the chromosome. As illustrated by the annotations from MEGABASE shown in the figure, the four loci are composed of chromatin of alternating types — L1 and L3 composed of type A chromatin, and L2 and L4 composed of type B. (C and D) The cumulative distribution functions show that loci composed of chromatin belonging to the same type tend to be closer in space than otherwise, despite the interlaced order and despite lying at greater genomic distances. This phenomenon is observed in FISH experiments and it is correctly predicted by our ChIP-Seq based modeling. The comparison between the predicted and the measured probability distributions shows excellent agreement for both the average distance and the distance fluctuations. (See SI for more examples of validation with FISH data).

Representative predicted 3D conformations for chromosome 2 and for chromosome 10 are shown in Figure 2. As previously observed in (15), the analysis of the conformational ensembles shows the existence of micro-phase separation between chromatin of different types leading to the formation of the characteristic patterns of interactions seen in Hi-C maps. Examples of the long-range patterns that are captured by our predictions are shown in Figure 2. The more transcriptionally active segments of chromatin (in compartments A1 and A2) are more frequently found on the outer surface while the inactive segments (in B1, B2, and B3) typically reside in the core of chromosomes.

The quality of the structural predictions achieved using the chromatin annotation inferred by MEGABASE shows that there exists a clear sequence-to-structure relationship between the sequences of chromatin types predicted from epigenetic marks and genome architecture. The accuracy achieved by using our energy landscape model in predicting the effects of compartmentalization, as seen by Hi-C and by 3D FISH, supports the plausibility of microphase separation being the physical process driving compartmentalization in chromosomes (15, 27-29) (Figure 4).

**Figure 4.**
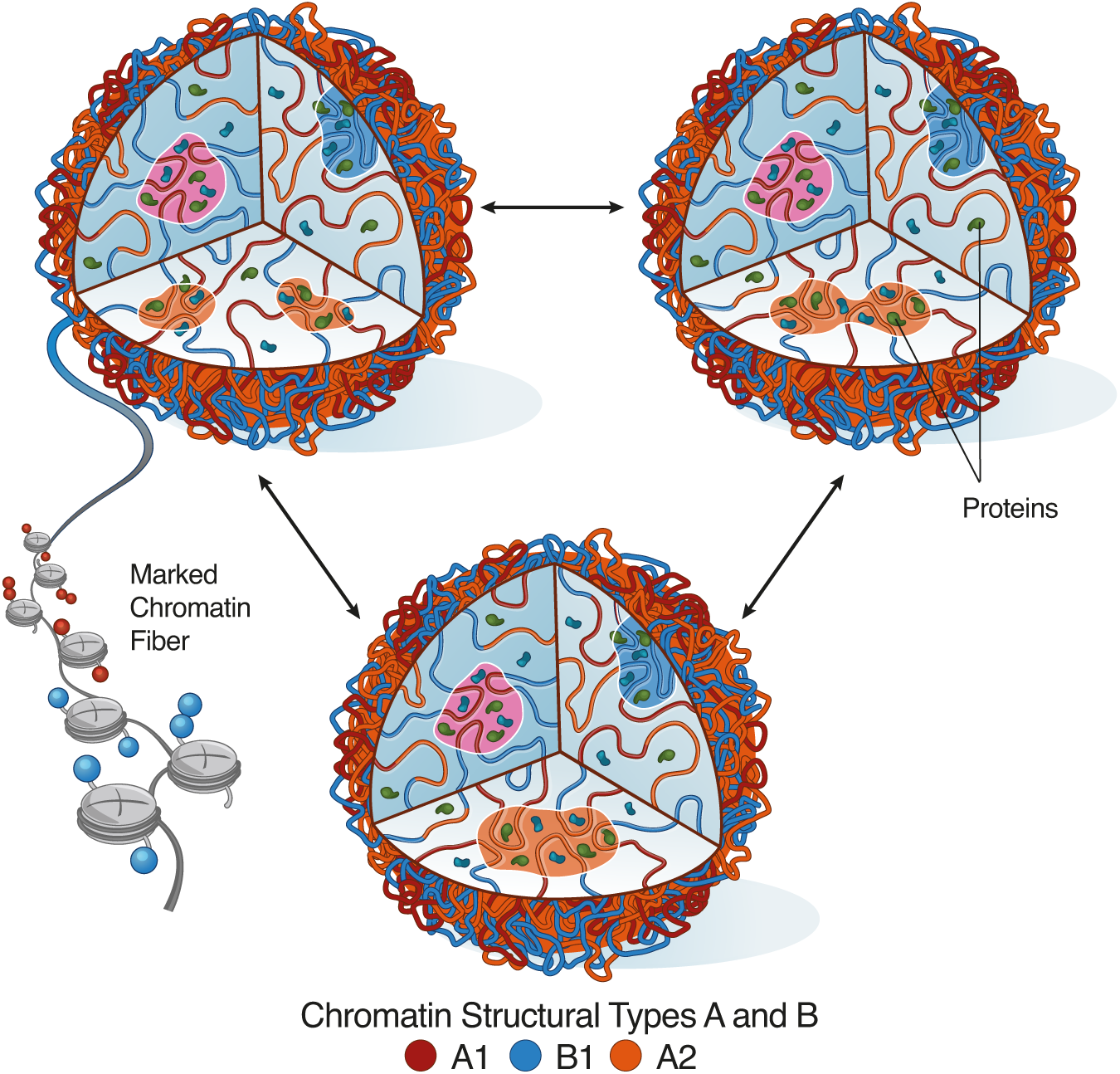
A process of micro phase separation explains compartmentalization in chromosomes. The Minimal Chromatin Model hypothesizes that chromatin characterized by homogeneous epigenetic markings undergoes a process similar to phase separation under the action of the proteome present in the nucleus. In simulations, we observe that segments of chromatin belonging to the same structural type tend to segregate, forming liquid droplets, which rearrange dynamically by splitting and fusing. This simple process of phase separation is sufficient to explain the emergence of compartmentalization in genomes as observed in DNA-DNA ligation assays and microscopy experiments.

The success achieved in reliably predicting chromosome architecture indicates that our probabilistic model captures the essential features of epigenetic marks that are associated with compartmentalization. Hence, we further exploit MEGABASE to study this relationship by calculating the content of mutual information shared between markers and compartments, and so quantifying which of the markers are the best predictors of compartmentalization. It is immediately evident that certain biochemical markers share a high content of mutual information with chromatin structural types while others do not. According to our neural network, histone methylations HK36me3, H3K27me3, H3K4me1, and H4K20me1 and nuclear proteins EED, ZBED1, TRIM22, and HCFC1 carry most of the information associated with identifying the chromatin types (Figure S27). In contrast, we see that although compartment A, for example, has a very high content of H3K27ac, that marker by itself is a poor predictor owing to its modest mutual information value.

Histone modifications alone carry enough information to predict genome architecture. To illustrate the disproportionate predictive value of histone marks, we created a reduced model by training MEGABASE using only the 11 patterns of histone modifications available in the ENCODE database. The sequences of chromatin types predicted by this reduced model turn out to be only marginally different from those obtained by the full data set of ChIP-Seq tracks (See SI).

Our results demonstrate clearly that it is possible to generate *de novo* predictions of the genome’s 3D structure, as well as specific predictions about the results of Hi-C and FISH experiments, using only ChIP-Seq data on histone modifications as an input. The faithfulness of the predicted conformational ensembles underlines the existence of a sequence-to-structure relationship between patterns of histone modifications and the 3D spatial arrangement of chromosomes.

These findings offer great hope that, like the problem of protein folding before it, the puzzle of genome folding may be amenable to computational predictions (30). Yet, despite the success of the neural network based prediction algorithm, the details of the mechanism underlying chromatin folding remain unclear. Does chromatin fold into a specific conformation because of the particular sequence of epigenetic markers, or, vice versa, do compartments share similar epigenetic markers because of chromosome architecture? Dynamical studies using Hi-C and other methods will doubtless be essential in addressing these questions.

## Data Availability

All simulated Hi-C contact maps in this manuscript can be viewed interactively using Juicebox (http://www.aidenlab.org/juicebox/), and directly at http://bit.ly/2vcBrVF.

## Footnotes

1 The average ratio between simulated distances and FISH-measured distances has been used to calibrate the length scale of simulation. One unit of length in simulation resulted to correspond to length of 6.05*μm*, which also implies the size of a simulated chromosomal territory being approximately 2-3 μm across— consistent with what was previously reported in ref. 2. Cremer T & Cremer C (2001) Chromosome territories, nuclear architecture and gene regulation in mammalian cells. *Nat Rev Genet* 2(4):292-301.

## References

1. Bickmore WA (2013) The Spatial Organization of the Human Genome. Annual Review of Genomics and Human Genetics, Vol 14 14:67–84.

2. Cremer T & Cremer C (2001) Chromosome territories, nuclear architecture and gene regulation in mammalian cells. Nat Rev Genet 2(4):292–301.

3. Whalen S, Truty RM, & Pollard KS (2016) Enhancer-promoter interactions are encoded by complex genomic signatures on looping chromatin. Nat Genet 48(5):488–496.

4. Krijger PH, et al. (2016) Cell-of-Origin-Specific 3D Genome Structure Acquired during Somatic Cell Reprogramming. Cell Stem Cell 18(5):597–610.

5. Rao SSP, et al. (2014) A 3D Map of the Human Genome at Kilobase Resolution Reveals Principles of Chromatin Looping. Cell 159(7): 1665–1680.

6. Gondor A (2013) Dynamic chromatin loops bridge health and disease in the nuclear landscape. Semin Cancer Biol 23(2):90–98.

7. Krijger PH & de Laat W (2016) Regulation of disease-associated gene expression in the 3D genome. Nat Rev Mol Cell Biol 17(12):771–782.

8. Fullwood MJ, et al. (2009) An oestrogen-receptor-alpha-bound human chromatin interactome. Nature 462(7269):58–64.

9. Montefiori L, et al. (2016) Extremely Long-Range Chromatin Loops Link Topological Domains to Facilitate a Diverse Antibody Repertoire. Cell Rep 14(4):896–906.

10. Lieberman-Aiden E, et al. (2009) Comprehensive Mapping of Long-Range Interactions Reveals Folding Principles of the Human Genome. Science 326(5950):289–293.

11. Dixon JR, et al. (2012) Topological domains in mammalian genomes identified by analysis of chromatin interactions. Nature 485(7398):376–380.

12. Eagen KP, Hartl TA, & Kornberg RD (2015) Stable Chromosome Condensation Revealed by Chromosome Conformation Capture. Cell 163(4):934–946.

13. Sexton T, et al. (2012) Three-Dimensional Folding and Functional Organization Principles of the Drosophila Genome. Cell 148(3):458–472.

14. Wang S, et al. (2016) Spatial organization of chromatin domains and compartments in single chromosomes. Science 353(6299):598–602.

15. Di Pierro M, Zhang B, Aiden EL, Wolynes PG, & Onuchic JN (2016) Transferable model for chromosome architecture. P Natl Acad Sci USA 113(43):12168–12173.

16. Zhang B & Wolynes PG (2016) Shape Transitions and Chiral Symmetry Breaking in the Energy Landscape of the Mitotic Chromosome. Phys Rev Lett 116(24): 248101.

17. Zhang B & Wolynes PG (2015) Topology, structures, and energy landscapes of human chromosomes. P Natl Acad Sci USA 112(19):6062–6067.

18. Grigoryev SA, et al. (2016) Hierarchical looping of zigzag nucleosome chains in metaphase chromosomes. P Natl Acad Sci USA 113(5):1238–1243.

19. Sanborn AL, et al. (2015) Chromatin extrusion explains key features of loop and domain formation in wild-type and engineered genomes. P Natl Acad Sci USA 112(47):E6456–E6465.

20. Phillips JE & Corces VG (2009) CTCF: Master Weaver of the Genome. Cell 137(7):1194–1211.

21. Nichols MH & Corces VG (2015) A CTCF Code for 3D Genome Architecture. Cell 162(4):702–705.

22. Zhang B & Wolynes PG (2017) Genomic Energy Landscapes. Biophys J 112(3):427–433.

23. Hopfield JJ (1982) Neural Networks and Physical Systems with Emergent Collective Computational Abilities. P Natl Acad Sci-Biol 79(8):2554–2558.

24. Lapedes A, Giraud B, & Jarzynski C (2002) Using Sequence Alignments to Predict Protein Structure and Stability With High Accuracy. Los Alamos National Laboratory Preprint.

25. Ekeberg M, Lovkvist C, Lan YH, Weigt M, & Aurell E (2013) Improved contact prediction in proteins: Using pseudolikelihoods to infer Potts models. Phys Rev E 87(1).

26. Bryngelson JD, Hopfield JJ, & Southard SN (1990) A protein structure predictor based on an energy model with learned parameters. Tetrahedron Computer Methodology 3(3): 129–141.

27. Hnisz D, Shrinivas K, Young RA, Chakraborty AK, & Sharp PA (2017) A Phase Separation Model for Transcriptional Control. Cell 169(1): 13–23.

28. Larson AG, et al. (2017) Liquid droplet formation by HP1alpha suggests a role for phase separation in heterochromatin. Nature 547(7662):236–240.

29. Strom AR, et al. (2017) Phase separation drives heterochromatin domain formation. Nature 547(7662):241–245.

30. Wolynes PG (2015) Evolution, energy landscapes and the paradoxes of protein folding. Biochimie 119:218–230.

